# Established *Pseudomonas syringae* pv. *tomato* infection disrupts immigration of leaf surface bacteria to the apoplast

**DOI:** 10.1101/2024.08.29.610363

**Authors:** Kimberly N. Cowles, Arjun S. Iyer, Iain McConnell, Ellie G. Guillemette, Dharshita Nellore, Sonia C. Zaacks, Jeri D. Barak

## Abstract

Bacterial disease alters the infection court creating new niches. The apoplast is an oasis from the hardships of the leaf surface and is generally inaccessible to nonpathogenic members of the phyllosphere bacterial community. Previously, we demonstrated that *Salmonella enterica* immigrants to the leaf surface can both enter the apoplast and replicate due to conditions created by an established *Xanthomonas hortorum* pv. *gardneri* (Xhg) infection. Here, we have expanded our investigation of how infection changes the host by examining the effects of another water-soaking pathogen, *Pseudomonas syringae* pv *tomato* (Pst), on immigrating bacteria. We discovered that, despite causing macroscopically similar symptoms as Xhg, Pst infection disrupts *S. enterica* colonization of the apoplast. To determine if these effects were broadly applicable to phyllosphere bacteria, we examined the fates of immigrant Xhg and Pst arriving on an infected leaf. We found that this effect is not specific to *S. enterica*, but that immigrating Xhg or Pst also struggled to fully join the infecting Pst population established in the apoplast. To identify the mechanisms underlying these results, we quantified macroscopic infection symptoms, examined stomata as a pinch point of bacterial entry, and characterized aspects of interbacterial competition. While it may be considered common knowledge that hosts are fundamentally altered following infection, the mechanisms that drive these changes remain poorly understood. Here, we investigated these pathogens to reach a deeper understanding of how infection alters a host from a rarely accessible, inhabitable environment to an obtainable, habitable niche.

**IMPORTANCE:** Pathogens dramatically alter the host during infection. Changes in host physical and biochemical characteristics benefit the pathogen and can reshape the composition of the bacterial community. In fact, rare members of the plant microbiota, namely bacterial human pathogens, such as *Salmonella enterica,* thrive in some plant infection courts. The increased success of human pathogens results from the conversion of the rarely accessible, inhabitable apoplast to an obtainable, habitable niche following infection. Here, we compared two phytopathogens, *Pseudomonas syringae* pv. *tomato* and *Xanthomonas hortorum* pv. *gardneri* within a tomato host and uncovered relevant niche changes potentially overlooked by the similarity in macroscopic symptoms. We investigated mechanisms used to reshape the host environment to the pathogen’s benefit and either success or failure of newly arriving immigrant bacteria. This study reveals information about bacterial disease of leaves and key changes that remodel inhospitable niches to new, conducive environments in the diseased host.

## Introduction

Hosts become a fundamentally altered niche following infection. Any plant pathology textbook offers many examples of how pathogens can modify the morphology of hosts with the level of modification ranging from intracellular to the whole organism (1). While these changes are well-documented, it remains unknown how changes to the host create new and preferred niches for organisms beyond the infecting pathogen. We and others have demonstrated that infected plants increase the incidence of rare members of the plant microbiota, namely bacterial human pathogens, such as *Salmonella enterica* (2–11). This increase in incidence and population growth results from the conversion of the rarely accessible, inhabitable interior space of the leaf, the apoplast, to an obtainable, habitable niche following infection.

Epiphytic bacteria, those found on the surface of plants, tolerate a harsh environment with rapidly fluctuating conditions. Survival of recent bacterial immigrants on a leaf surface is mostly due to luck on arrival near or in an oasis of nutrient and water availability, the base of glandular trichomes or in the grooves between cells (12–14). Flagellar motility and the capacity to form aggregates, either inter- or intra-species, increases an immigrant’s probability of survival (15, 16). Success requires adaptation to changes in water or nutrient availability, temperature, and UV irradiation levels, among others. In addition to abiotic factors, epiphytic bacteria must also contend with the plant immune response. Host surface receptors recognize conserved bacterial motifs and initiate a cascade of defenses meant to thwart potential invaders and control bacterial populations through resistance (for review (17)). Bacterial phytopathogens lacking cell wall degrading enzymes abandon the leaf surface and choose the apoplast for their infection court. The leaf apoplast has several obvious advantages over the leaf surface: protection from most UV irradiation, little to no cuticular wax encasing plant cell surfaces, and reduced fluctuation in free water. Although the existence of bacteria in the apoplast is well documented by microscopy (18), along with many bacterial factors that are necessary for disease, in general, little is known about the dynamic colonization of an infected apoplast during disease progression and distinct changes in the infected host.

One of most widely studied bacterial-plant interactions is *Arabidopsis thaliana* as a model host for *Pseudomonas syringae* pv. *tomato* (Pst). Pst uses a jasmonic acid mimic coronatine (COR) to open stomata for access to the leaf apoplast (19). Pst then further manipulates the host immune system using Type III effectors HopM1 and AvrE to induce the abscisic acid pathway and to close stomata to increase water potential in the apoplast (20–22). These findings are the foundational understanding of the disease Pst causes in tomato, bacterial speck. A macroscopically similar disease, bacterial spot of tomato, is caused by four lineages of *Xanthomonas*: *X. hortorum* pv. *gardneri* (hereafter referred to as Xhg), *X. euvesicatoria* pv. *euvesicatoria, X. euvesicatoria* pv. *perforans*, and *X. vesicatoria* (23–25). Tomato infection with either Pst or Xhg is characterized by water-soaked lesions and abundant phytobacterial growth. Multiple works from our lab have demonstrated that Xhg infection alters the tomato host in ways that benefit non-phytopathogenic bacteria inhabiting the leaf surface (2–5).

Although the primary purpose of altering the plant environment during infection is likely for the benefit of the pathogen, sweeping changes in physical and biochemical characteristics of the host reshape the composition of the bacterial community in an infection court as shown in human infections (26), but overlooked in plants. We have found that the dramatic change to the apoplast as a result of Xhg infection creates an available and habitable niche for bacteria that are usually precluded from stomatal entry and exiled to the leaf surface, such as *S. enterica* (2–5). Xhg infection does two things: 1) permits *S. enterica* access to the apoplast, a niche that the human pathogen can’t access on its own and 2) transforms the apoplast into a habitable niche for *S. enterica*, altering the apoplast in ways that promote bacterial replication (2–5). However, the mechanisms driving changes to the host that create new and preferred niches for organisms beyond the infecting pathogen and that influence bacterial dynamics in the apoplast remain unknown.

Here, we expanded our investigation of how leaf infection impacts the host and examined whether bacteria that create a water-soaked apoplast during infection, in general, permit leaf surface bacteria entry to this altered niche. We discovered that, unlike Xhg infection which promotes *S. enterica* success, Pst infection delays *S. enterica* colonization of the apoplast. In addition, we found that this effect is not specific to *S. enterica*, but that immigrating populations of Xhg or Pst also struggled to fully join the infecting Pst population established in the apoplast. These results support several possible mechanisms that we began to investigate in this work: 1) changes to a Pst-infected plant result in a barrier to apoplast entry, 2) Pst infection creates an inhospitable niche, or 3) macroscopically imperceptible distinctions between immigrating bacteria and established Pst populations impact bacterial survival. By comparing and contrasting strategies used by Pst and Xhg, this study provides fundamental information about bacterial disease of leaves and reveals mechanisms used to reshape the host environment to a new, conducive niche in the diseased host.

## Results

### Pst infection delays successful *S. enterica* colonization of tomato leaves

Previously, we had shown that an established Xhg infection promotes the growth of newly arriving *S. enterica* on tomato leaves (3). To determine if another phytopathogen that causes water-soaking, Pst, also enhances *S. enterica* persistence, we examined the impact of Pst infection on *S. enterica* populations over time. As done previously, UV irradiation was used to distinguish surface localized *S. enterica* from *S. enterica* found within the apoplast (3). Here, UV treatment was not a significant factor in bacterial populations (*P* > 0.01), and data from UV-treated and non-UV-treated samples were collapsed to create Figure 1. At the arrival site (Fig. 1A), *S. enterica* suspensions were absorbed equally amongst the three treatments, and no significant differences in *S. enterica* populations were observed at 3 hours post-arrival (HPA) between treatments or from the starting inoculum. Xhg infection resulted in higher apoplastic *S. enterica* populations compared to the initial arriving population and compared to plants treated with Pst or water infiltration at 24, 48, and 72 HPA. Contrastingly, *S. enterica* populations didn’t significantly increase in Pst-infected leaves until 48 HPA and remained lower than *S. enterica* populations in Xhg-infected leaves through 72 HPA. Similar to Xhg infection, water congestion in healthy leaves also resulted in increased *S. enterica* populations compared to the initial arriving population starting at 24 HPA. However, as with Pst-infected leaves, water congested healthy leaves supported lower overall levels of *S. enterica* compared to Xhg-infected leaves.

**Figure 1.**
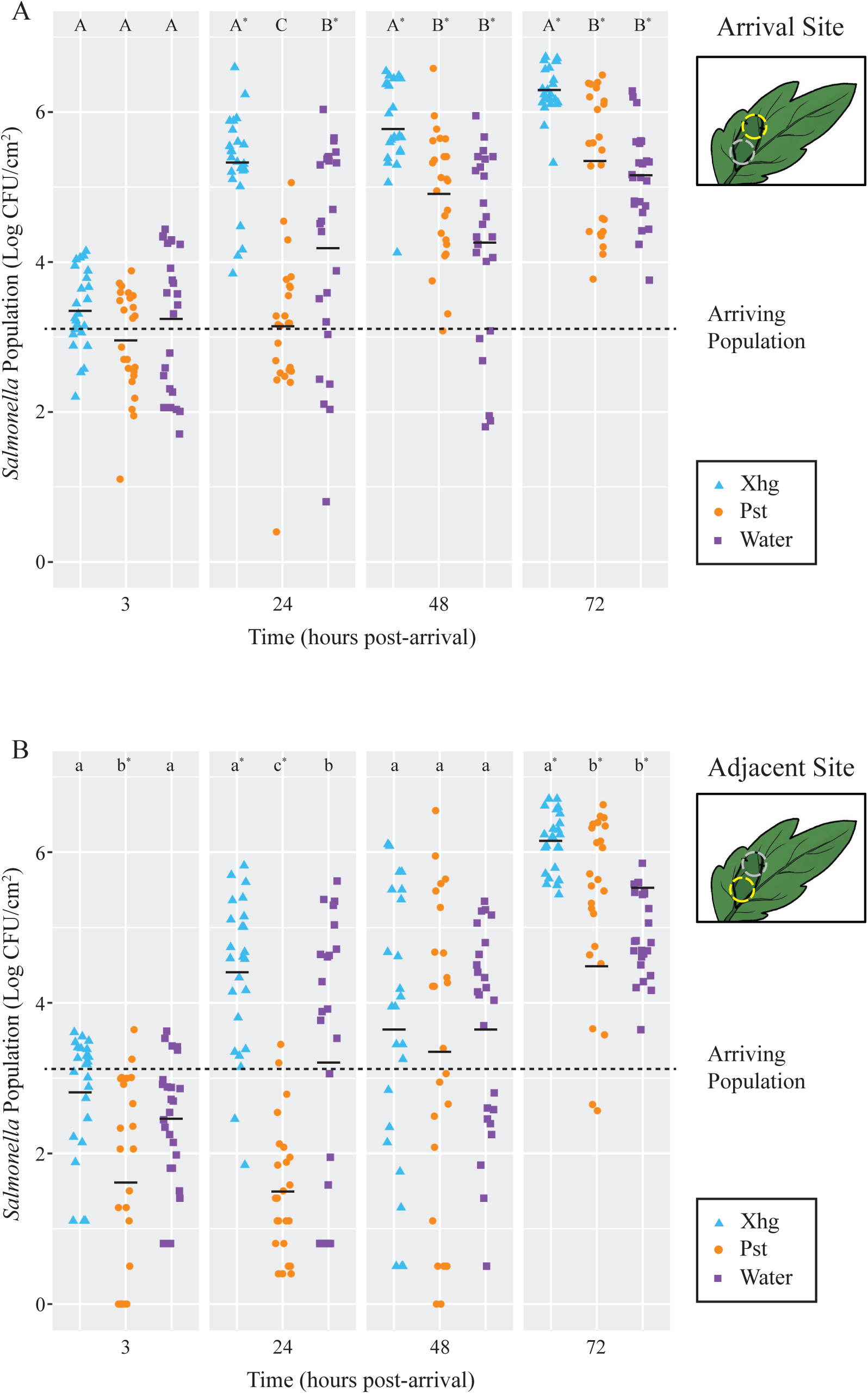
Immigrating *S. enterica* have a delayed benefit from established Pst infection. *S. enterica* populations were monitored 3, 24, 48, or 72 hours after arrival on tomato leaves previously infiltrated with Xhg (48 HPI; cyan triangles), Pst (48 HPI; orange circles), or water (0 HPI; purple squares). Leaves were sampled at the *S. enterica* arrival site (A) and a distinct adjacent site within the infiltrated area (B). Data from three independent experiments are presented as log CFU/cm^2^, and each symbol represents bacterial populations from one tomato leaf. Half of the leaves from each treatment and time point were treated with UV irradiation but data were collapsed as there was no significant difference between UV-treated and non-UV-treated samples (*P*>0.01). The dashed line indicates the arriving *S. enterica* population. Means for each treatment at each time point are depicted with horizontal black lines. Letters denote significant differences between treatments within a single time point and leaf site, and asterisks indicate significant differences from the initial arriving population (*P<*0.05). Combining three independent experiments, n=24 leaves per treatment per time point.

To examine migration of arriving bacteria, we monitored bacterial populations at an additional site within the infiltrated area, which we termed the adjacent site. Bacterial populations at the adjacent site showed similar patterns as the arrival site with several notable differences. First, unlike at the arrival site, *S. enterica* populations at the adjacent site in Pst-infected leaves are smaller compared to the arriving population at 3 and 24 HPA (Fig. 1B). Second, *S. enterica* populations at the adjacent site on healthy water congested leaves did not grow from arriving population levels until 72 HPA (Fig. 1B), instead of at 24 HPA as seen at the arrival site (Fig. 1A). Third, although these experiments tend to show a relatively large amount of variation between samples and individual plants, all treatments displayed a wider range in *S. enterica* population sizes at the adjacent site, when compared to the arrival site, at 48 HPA. This 6-log range of bacterial population size resulted in no significant differences amongst treatments or compared to the arriving population at this timepoint. Phytopathogen populations were not significantly different from one another at any time point at either the arrival site or the adjacent site (*P* > 0.05; Fig. S1).

### Xhg- and Pst-infected leaves have similar quantifiable disease symptoms

To identify differences in Xhg and Pst infection that could explain the delayed colonization of *S. enterica*, we characterized multiple quantifiable phenotypes from Xhg- and Pst-infected leaves. First, to simulate a natural infection, tomato plants were dip-inoculated with Xhg and Pst suspensions and resulting disease symptoms were photographed at 1-4 days post-inoculation (DPI). To quantify disease symptoms, we developed the Leaf Lesion Detector application to quantify lesion numbers, size, and the percent infection observed over time in Xhg- and Pst-infected leaves (Fig. 2). Representative images from 2 DPI leaves are shown at multiple stages of the application analysis. The Leaf Lesion Detector application segments images using a series of hue, saturation, value (HSV) thresholding operations, and contour finding (27) on greyscale-converted images, both of which use empirically predetermined, but configurable pixel-value limits. Each leaf and reference image is processed as follows: 1) Segment reference area by thresholding, applying noise reduction, then counting resulting pixels. (not shown) 2) Segment leaf area by thresholding and contour finding (Fig. 2A, Total Area and Outline). 3) Segment lesion area by thresholding, contour finding, labelling all lesions and summing the area for all lesions above the configured minimum lesion size threshold pixel count (Fig. 2A, Lesions). All the segmentations are combined and a color map applied to the lesions by size to create the final image (Fig. 2A, Modified). If the total lesion area of a leaf exceeds 3.5 %, the image is reprocessed with a more stringent lower bound (supplied in the configuration) for lesion size. After testing several different cutoff values on a range of test images, 3.5% was chosen as it appeared to give the best trade-off between maximizing detection and minimizing false positives. The percent infected area is calculated as a ratio of lesion and leaf pixel counts. Conversion from pixel count to mm^2^ is handled by multiplying a segment’s pixel count by the ratio of known reference area to known reference pixel count. No significant differences in infection characteristics were measured over the four day time course using the Leaf Lesion Detector app (Fig. 2B).

**Figure 2.**
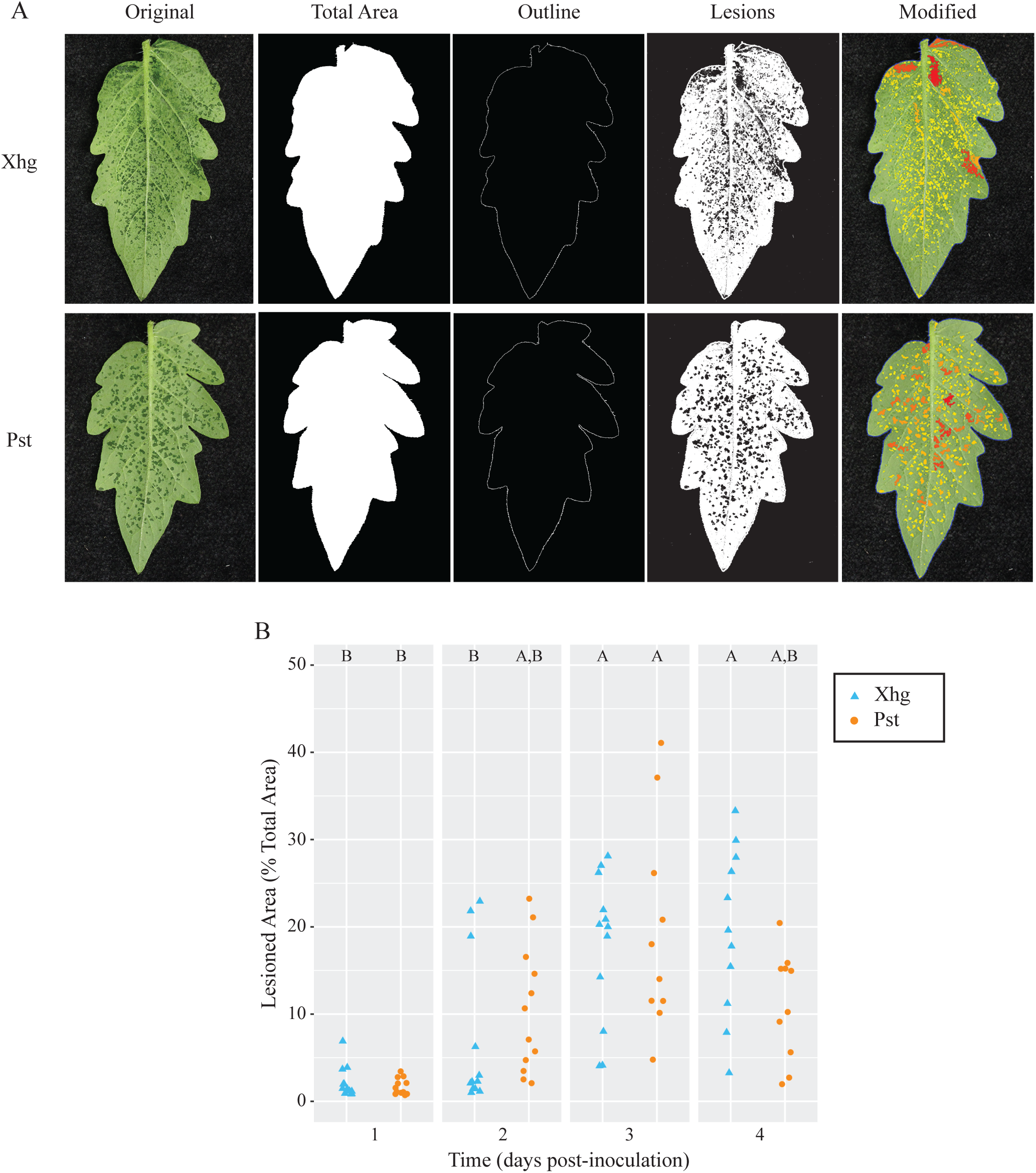
Water-soaking symptoms in Xhg- and Pst-infected leaves are macroscopically similar. (A) Representative images from Xhg- and Pst-infected leaves at 2 DPI are shown at multiple stages of the Leaf Lesion Detector application analysis. Each original leaf image is processed to identify total area, leaf outline, and lesions. The modified image is presented as a color map based on lesion size. (B) Lesions are quantified as the percentage of the total leaf area of Xhg-infected (cyan triangles) or Pst-infected (orange circles) leaves at 1-4 DPI. Letters denote significant differences between treatments within a single time point (*P<*0.05). Combining three independent experiments, n=12 leaves per treatment per time point.

### Xhg- and Pst-infected leaves have similar macroscopic and microscopic symptoms

*S. enterica* persistence in Figure 1 was quantified in infiltrated plants. To qualitatively examine plant disease under those conditions, infiltrated plants were imaged over time, and representative images from infection of each pathogen are shown in Fig. 3A. No qualitative differences were observed when comparing infected leaves at each time point. Both Xhg- and Pst-infiltrated leaves showed patchy water soaking at 1DPI, complete water soaking of the infiltrated area by 2 DPI, a combination of water soaking and the beginnings of necrosis at 3 DPI, and drier, more necrotic tissue by 4 DPI (Fig. 3A).

**Figure 3.**
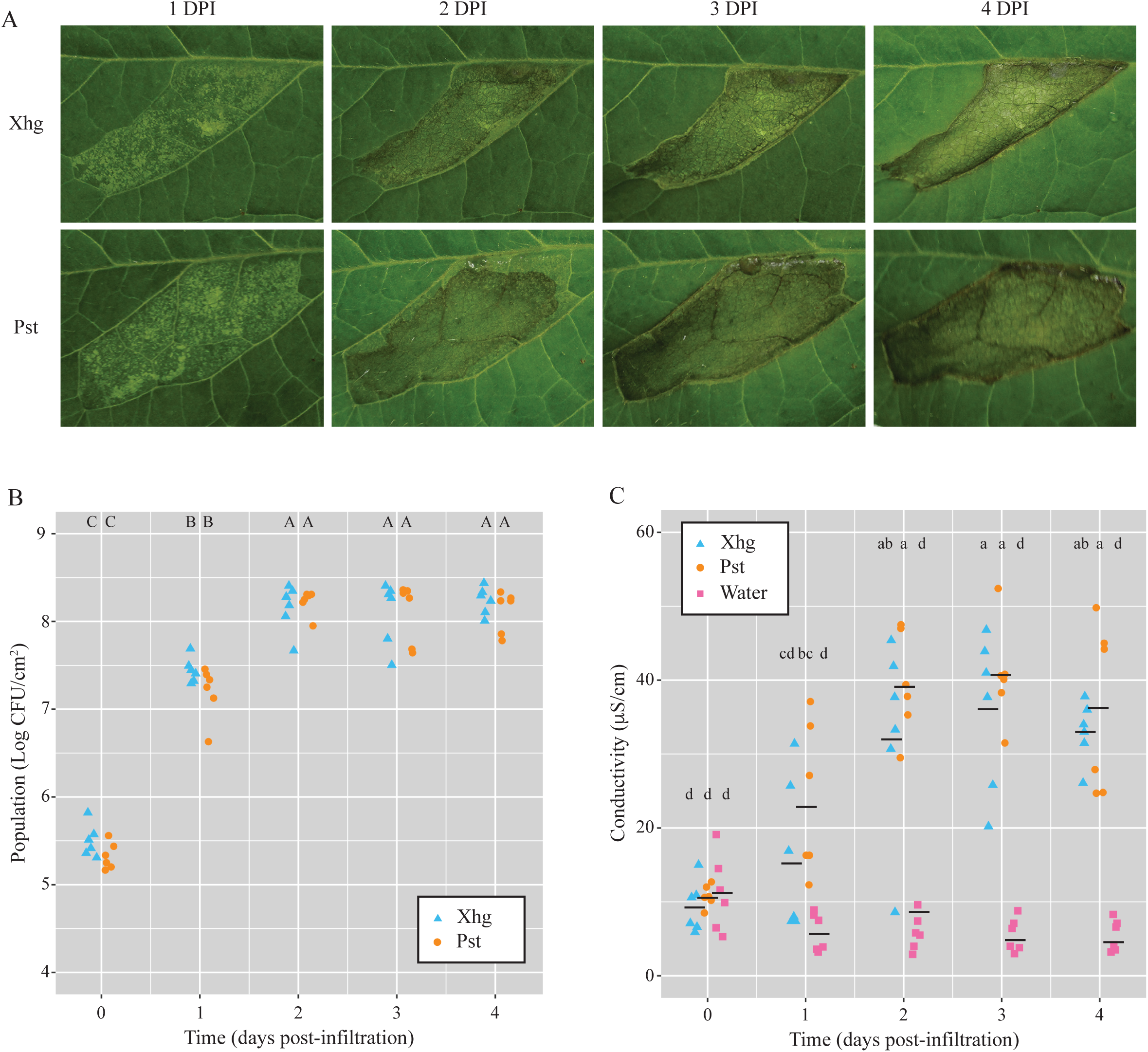
Leaves infiltrated with Xhg or Pst have indistinguishable symptomology, bacterial populations, and cellular damage. (A) Representative images from Xhg- and Pst-infiltrated leaves from 1-4 DPI. Images were taken of the same leaves on consecutive days. (B) Bacterial populations were monitored 0-4 DPI from tomato leaves infiltrated with Xhg (cyan triangles) or Pst (orange circles). Leaves were sampled within the infiltrated areas, and Xhg and Pst populations are presented as log CFU/cm^2^. (C) Electrolyte leakage was measured by conductivity levels (μS/cm) from 0-4 DPI in leaves infiltrated with Xhg (cyan triangles), Pst (orange circles), or water (pink squares). Each symbol represents bacterial populations or conductivity readings from one tomato leaf. Means for each treatment at each time point are depicted with horizontal black lines. Letters denote significant differences between treatments over time (*P<*0.05). Combining three independent experiments, n=6 leaves per treatment per time point.

In addition to visible symptoms following infection, we measured both bacterial populations and cellular damage in infiltrated areas. As with symptomology, no differences were detected between Xhg- and Pst-infected leaves. Infiltrated pathogens reached a carrying capacity of 8.0-8.5 Log CFU/cm^2^ by 2 DPI, and bacterial populations were statistically equivalent at each time point examined (Fig. 3B). Electrolyte leakage measured by conductivity was used as a proxy for cellular damage (28) throughout disease progression compared to leaves infiltrated with water. In parallel with lesion development, infection with both pathogens resulted in increasing levels of conductivity, and no differences were detected between Xhg- and Pst-infected leaves (Fig. 3C).

### Stomatal aperture patterns differ between Xhg- and Pst-infected leaves

Previous work has shown that Pst opens and closes stomata of *Arabidopsis thaliana* or tomato leaves depending on the stage of infection (21, 22). To monitor stomatal aperture patterns, we infiltrated leaves with phytopathogen or water and used a dental resin method to capture impressions of stomata over time. Stomatal apertures were measured using ImageJ and indicated by a ratio of width to length. The outline of stomata results in some measurement value for both length and width, making it impossible to get a ratio of 0.0. Thus, we considered a stomatal aperture ratio of 0.25 to be closed. Larger stomatal aperture ratios indicate more “open” stomata. Analysis of the resulting impression images demonstrated that while some differences exist between Xhg- and Pst-infected leaves, infection with either pathogen results in open stomata at 48 HPI (Fig. 4A), the point in time when *S. enterica* cells were applied in our earlier experiments (Fig. 1). All treatments had relatively closed stomata at 1 and 4 HPI (Fig. 4A). Pst-infected leaves had open stomata by 24 HPI while Xhg-infected leaves had open stomata at 48 and 72 HPI (Fig. 4A). Water infiltration resulted in open stomata at 24 and 48 HPI, but stomata were closed at 72 HPI (Fig. 4A).

**Figure 4.**
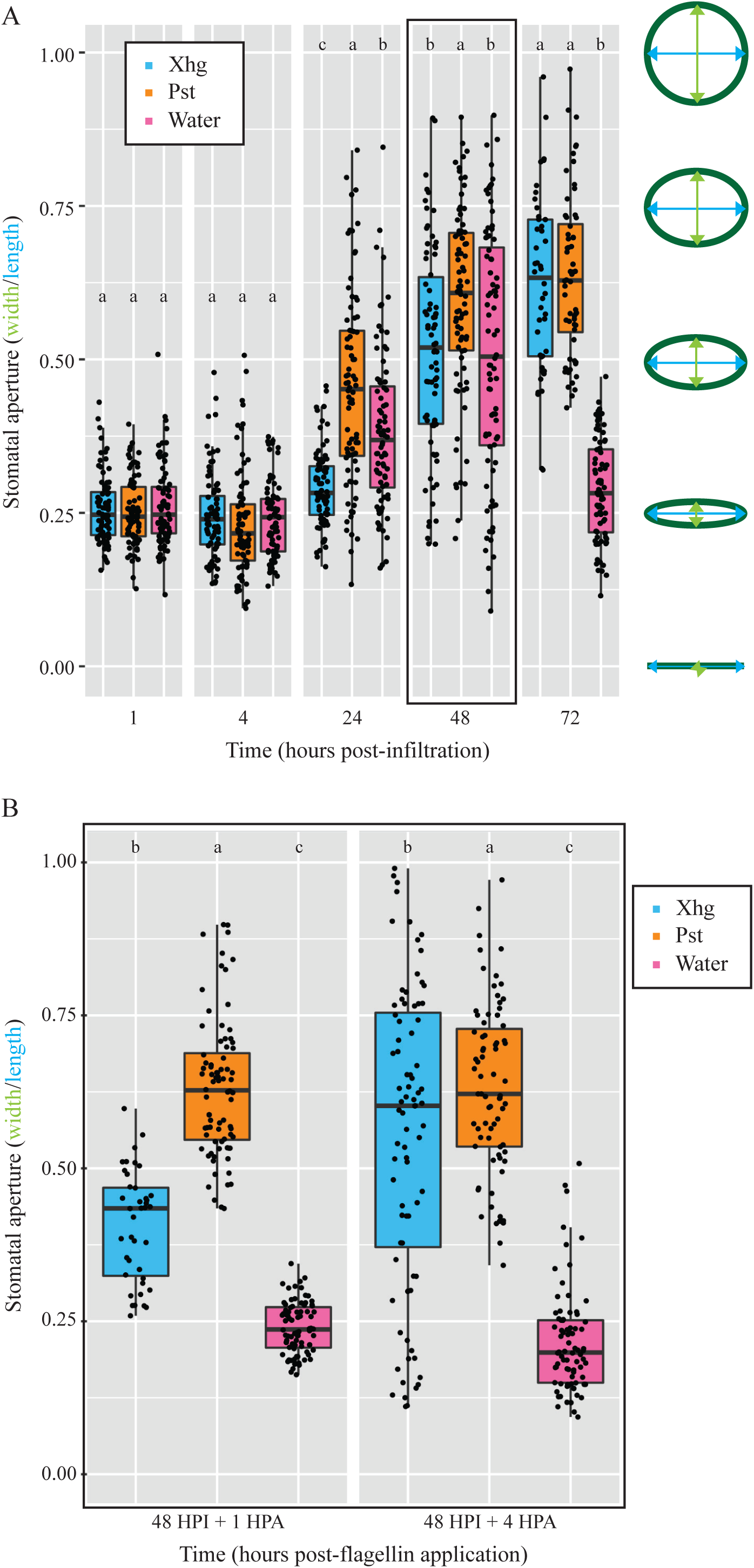
Both Xhg- and Pst-infiltrated leaves have open stomata upon arrival of immigrating *S. enterica* at 48 HPI. (A) Stomatal apertures were monitored at 1, 4, 24, 48, and 72 HPI in tomato leaves infiltrated with Xhg (cyan), Pst (orange), or water (pink). Apertures were measured using ImageJ and indicated by a ratio of width (green) to length (blue) as depicted on the right. The 48 HPI data are outlined in black to highlight the point when *S. enterica* arrives at the infected area in other experiments. (B) Stomatal apertures were measured at 1 and 4 HPA of flagellin from leaves that had been previously infiltrated with Xhg (cyan), Pst (orange), or water (pink) 48 hours earlier (48 HPI). Data from three independent replicates of each experiment are represented as boxplots with each symbol corresponding to one stomatal aperture (n>50 per treatment per time point). Letters denote significant differences between treatments within a single time point (*P<*0.05).

In our *S. enterica* persistence experiments (Fig. 1), both *S. enterica* and either phytopathogen were present on leaves at the same time while the stomatal aperture experiments in Fig. 4A did not have *S. enterica*. We hypothesized that stomatal aperture regulation could be influenced by *S. enterica,* and we repeated the stomata experiments with the addition of *S. enterica* flagellin, a known signal for stomatal aperture movement (20). As done above, leaves were infiltrated with a phytobacterial pathogen or water, and infection was allowed to proceed for 48 HPI. Then, flagellin was spotted on the surface of infiltrated areas, similar to *S. enterica* arrival in persistence experiments, and resin impressions were taken 1 or 4 hours later (48 HPI + 1 and 48 HPI + 4). In persistence experiments (Fig. 1), droplets of *S. enterica* suspensions were absorbed into leaves by 3-4 hours post-arrival, so these time points represent stomatal apertures present at the time of *S. enterica* arrival and absorption. At 48 HPI, water-treated leaves had open stomata in the absence of flagellin (Fig. 4A) and closed stomata in the presence of flagellin (Fig. 4B). In contrast, Pst-infected leaves had comparably open stomata in both the presence and absence of flagellin (Fig. 4A-B). Xhg-infected leaves had closed stomata immediately upon flagellin application, but apertures returned to a more open state after 4 hours (Fig. 4B).

### Phytopathogens target potential competing bacteria in an *in vitro* killing assay

Our data demonstrate that Xhg infection creates a conducive environment for *S. enterica* persistence (2–5) (Fig. 1). To test the hypothesis that Pst can be hostile towards other bacteria, we performed ‘killing assays’ used to study bacterial Type VI secretion systems (29). Two type VI secretion systems are present in the Pst genome (CP034558 (30)) while Xhg does not have the genes needed for Type VI secretion(31), suggesting a potential mode of action for the differential effects on *S. enterica* survival. These data demonstrate that incubation with homogenized healthy leaf tissue has no impact on *S. enterica* levels while infected tissue containing Xhg or Pst negatively affected *S. enterica* (Fig. 5A). Incubation with Xhg-infected tissue reduces *S. enterica* populations by ∼0.25 log while incubation with Pst-infected tissue results in ∼1.25 log reduction in *S. enterica* (Fig. 5A).

**Figure 5.**
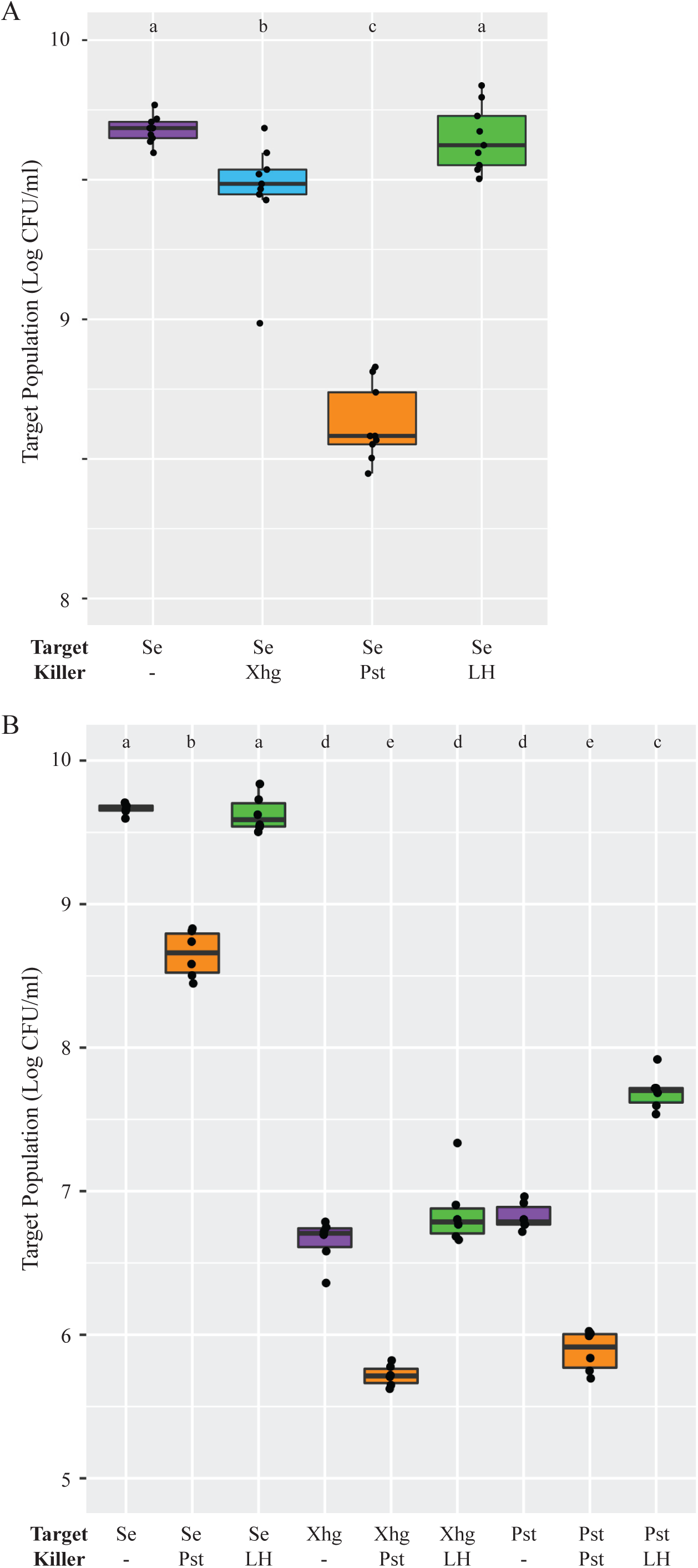
*In planta* grown Pst inhibits *in vitro* grown *S. enterica.* (A) Using a ‘killing assay’ for Type VI secretion activity, *S. enterica* populations were measured after 24 hour incubation with water (purple), *in planta* grown Xhg (cyan), *in planta* grown Pst (orange), or healthy leaf homogenate (green). (B) Type VI secretion ‘killing’ assays were repeated with three target bacterial strains (*S. enterica,* Xhg, and Pst) and three treatment conditions (water, purple; *in planta* grown Pst, orange; healthy leaf homogenate, green). Target bacterial population data from two independent experiments are presented as log CFU/ml in boxplots. Letters denote significant differences between treatments (*P<*0.05) and n=6.

To determine if *in planta* grown Pst affects other target bacteria, we performed additional ‘killing assays’ with *in vitro* grown *S. enterica,* Xhg, and Pst as the target strains. As seen previously, Pst-infected tissue lowered *S. enterica* populations by ∼1.25 log while homogenized tissues from healthy plants had no effect (Fig. 5B). As with *S. enterica*, incubation of Pst-infected tissue with Xhg or Pst reduced target populations by ∼ 1 log (Fig. 5B). While incubation with homogenized healthy leaf tissue had no impact on Xhg populations, Pst target populations increased ∼ 1 log when compared to incubation with water (Fig. 5B).

### A naïve, water-congested environment promotes phytopathogen success over pre-colonized tissues

With the knowledge that *in planta* grown Pst inhibits *S. enterica,* Xhg, and Pst growth, we monitored the fate of newly arriving Xhg or Pst (immigrants) to Xhg-or Pst-infected tissue, pre-colonized by “residents.” One day after arrival on water-congested, healthy leaf tissue, both Xhg (Fig. 6A-B) and Pst (Fig. 6C-D) immigrant populations were significantly larger compared to the arriving population, suggesting growth in healthy water-congested apoplasts that lacked resident bacterial populations. UV irradiation to remove surface bacteria (3) reduced both Xhg and Pst populations compared to non-UV treated samples in water-congested leaves. However, UV-treated samples, at the arrival site, still had higher populations when compared to the initial arrival population, suggesting that the phytobacterial pathogen immigrants migrated to the UV-protected apoplast and grew (Fig. 6A, C). UV-treated samples from the adjacent site were not significantly different from the arriving population (Fig. 6B, D).

**Figure 6.**
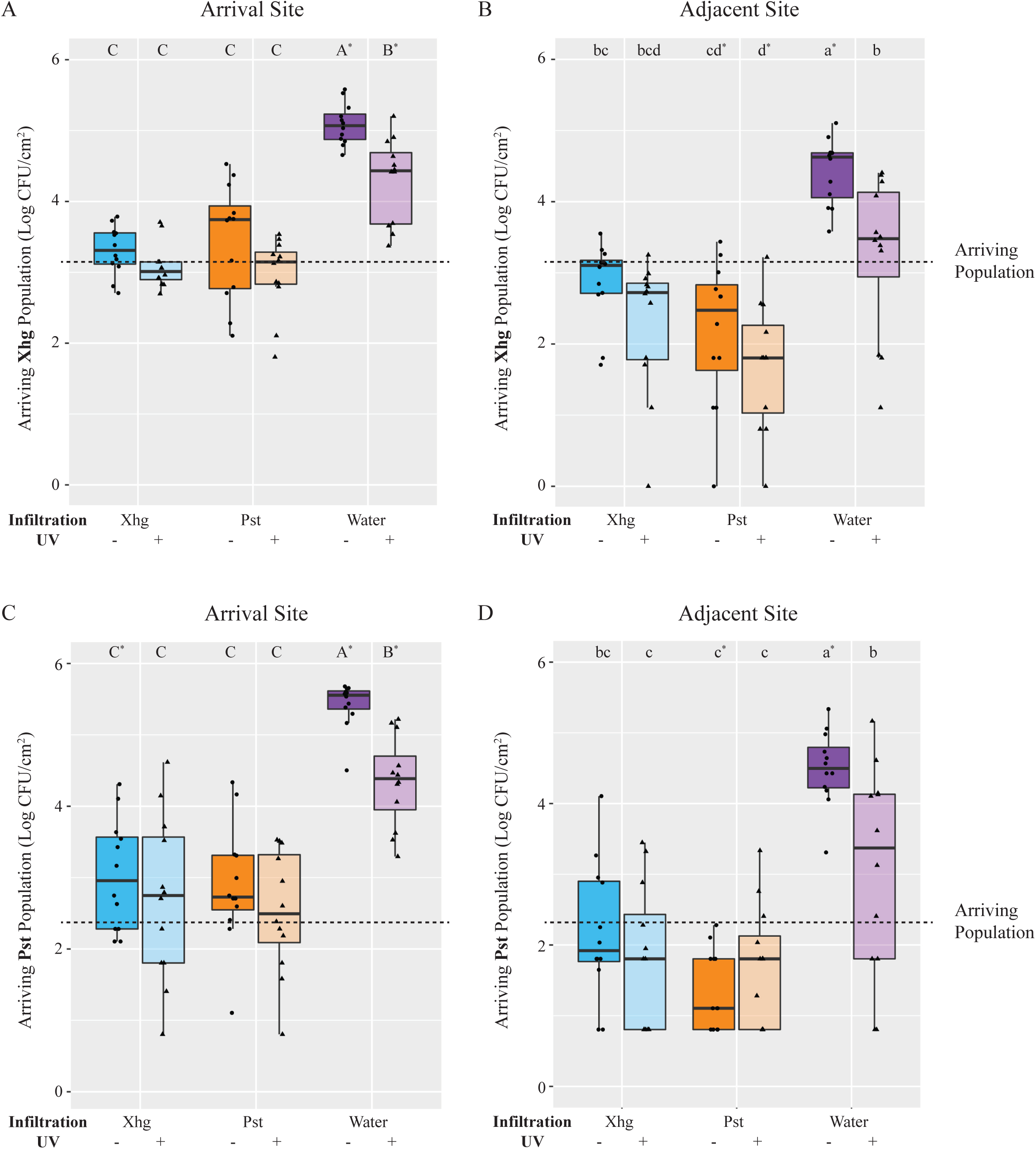
The fate of immigrating bacteria depends on the identity of the established infection. Xhg (A-B) and Pst (C-D) populations were monitored 24 hours after arrival on tomato leaves previously infiltrated with Xhg (48 HPI; cyan), Pst (48 HPI; orange), or water (0 HPI; purple). Leaves were sampled at the arrival site (A, C) and a distinct adjacent site within the infiltrated area (B, D). Data from three independent experiments are presented as log CFU/cm^2^ in boxplots, and each symbol represents bacterial populations from one tomato leaf (n= 12 leaves per treatment per time point). Half of the leaves from each treatment and time point were treated with UV irradiation and distinguished here with color shading. The dashed line indicates the arriving bacterial population. Letters denote significant differences between treatments within a single leaf site for each bacterium separately, and asterisks indicate significant differences from the initial arriving population (*P<*0.05).

In contrast to increased immigrant populations on water-congested, healthy leaves, leaves that were pre-colonized with resident Xhg or Pst, for the most part, either inhibited or reduced immigrant bacterial populations. In addition, unlike populations on water-congested, healthy leaves, UV irradiation had no impact on immigrant Xhg and Pst populations on infected leaf tissue (Fig. 6), most likely because the bulk of these bacteria had migrated into the UV-protected apoplast. Regardless of UV treatment, Xhg immigrants arriving on Xhg-infected leaves stayed at the same level or were reduced compared to the initial arriving populations (Fig. 6A-B). Contrastingly, Pst immigrants arriving on Xhg-infected leaves were either the same as the arriving population or slightly higher (Fig. 6C-D) but well below levels seen in water congested, healthy tissue, suggesting that Pst immigrants have some success on Xhg-infected leaves but replicate to greater numbers on naïve tissue. Commonly, the Pst-infected environment had a more negative impact on immigrant bacteria, especially *S. enterica* (Fig. 1) and Xhg (Fig. 6A-B), then the Xhg-infected leaf. For example, one day after arrival on Pst-infected leaves, Xhg immigrant populations were significantly reduced compared to the initial arriving populations (Fig. 6A-B). Comparably, one day after arrival on Pst-infected leaves, Pst immigrant populations either remained at arriving population levels or were reduced (Fig. C-D). In general, bacterial populations at the adjacent site showed similar patterns as those at the arrival site but had overall lower levels of recovered bacteria (Fig. 6).

## Discussion

The leaf surface is a highly heterogenous environment and can be characterized as a fragmented habitat driving the development of microbial communities (32–34). Factors that determine the maximum number of individuals on the leaf surface are host-driven (plant species, water pooling, or nutrient leaching through the cuticle) or bacteria-driven (environmental stress response, motility, or aggregate formation). An immigrant’s fate on the leaf surface is determined by the luck of arriving at or near an oasis (35) or with others (12, 36). One avenue used by phytopathogenic bacteria to infect leaves is entry to the leaf interior through stomata, the gas exchange portal. Early in infection, both Xhg and Pst transform the air-filled apoplast to an aqueous environment (reviewed in (37)). Based on our results, bacteria that arrive on a leaf as immigrants and attempt to establish themselves in the apoplast appear to encounter fundamentally different niches upon arrival to a naive host compared with a plant hosting a previously established infecting population. Furthermore, our data show that the identity of the resident population also influences the fitness outcomes for the immigrating bacteria.

### Nutrient Availability in the Apoplast

Upon arrival to a healthy, water-congested apoplast, all three examined bacteria, whether phytopathogenic or not, demonstrated a two-log increase in growth from the initial inoculum population (Figs. 1 and 6). The similar fate for bacteria on a healthy, yet water-congested, host suggests that uptake of the droplet containing bacteria concurrently allows bacterial entry to the apoplast. In addition, bacteria that reach the apoplast have access to some degree of available nutrients, even *S. enterica* which does not manipulate the host niche. A ‘dry’ healthy apoplast is mainly air-filled with a thin water film over cell surfaces. Apoplastic nutrients are thought to be in a bound state, attached to cell surfaces, or within surrounding cells, and relatively unavailable to invading bacteria (38, 39). It is hypothesized that an influx of water in the ‘wet’ healthy apoplast may disrupt metabolite uptake and result in a dysregulation of metabolite partitioning with higher than normal water-soluble metabolite concentrations in the apoplast (38, 40). This access to nutrients could lead to the observed increases in *S. enterica,* Xhg, and Pst growth (Figs. 1 and 6). The temporary relief from nutrient restriction appears to be nondiscriminatory and available to any bacterium lucky enough to arrive in a water droplet on the water-congested leaf.

Unlike the transient nature of abiotic water congestion described above, pathogen-induced water soaking is maintained over days and represents a more complex system of pathogen manipulation of the host and the host response. As with abiotically water-congested leaves, *S. enterica* replicates within the protected niche of the Xhg-infected apoplast within the first 24 hours of arrival on infected tissue (Fig. 1). Contrastingly, *S. enterica* doesn’t demonstrate growth in Pst-infected tissue until 48 HPI and doesn’t reach the same level as *S. enterica* populations in Xhg-infected tissue at any time point (Fig. 1). There are also differences in the fates of immigrant phytopathogens on infected tissue, compared with abiotically water-congested leaves. Neither Xhg nor Pst immigrants increase from arriving population levels on Xhg- or Pst-infected leaves (Fig. 6). The static nature of these populations could represent lack of replication, slow replication (that is not detectable within 24 HPI), or an equal rate of replication to death, resulting in no net increase in population levels. We hypothesize that the differential impact of resident phytopathogens and/or infections on immigrant success results from differences in available resources within the apoplast. Monopolization of resources within the established infection court by resident bacteria could lead to direct competition for space and/or nutrients.

The formation of distinct niches based on the identity of the infecting phytopathogen may be the result of the available nutrient profile. As mentioned above, water congestion itself leads to a misregulation of metabolite partitioning (38, 40) and growth of *S. enterica* (Fig. 1). Water soaking due to infection can also influence plant metabolism directly to the benefit of the resident organism. Phytobacterial pathogens such as Pst and *Ralstonia solanacearum* use secreted effectors to manipulate their host to create a more nutritionally favorable environment or increase specific nutrient availability (41, 42). Multiple additional phytopathogens also hijack plant metabolism to shift source leaves into sink leaves, providing the pathogens with access to nutrients (43–46). Alternatively, established Xhg populations may have distinct nutritional requirements from *S. enterica,* allowing both bacteria to thrive, while established Pst populations have overlapping nutritional needs, resulting in competition for resources. Future experiments examining the metabolite profiles of the infected apoplasts could reveal details of this mechanism.

### Stomatal Response and Bacterial Entry

Without cell wall degrading enzymes, bacteria like *S. enterica* have restricted access to the leaf interior through natural openings such as stomata, hydathodes, and wounds (47, 48). Although the primary function of stomata is the regulation of gas exchange and water retention, stomatal cells (guard cells) can recognize conserved microbe associated molecular patterns (MAMPs) and close the opening to reduce or prevent bacterial entry (19, 20, 39, 49). We see this closure response in our experiments when *S. enterica* flagellin is added to the surface of tissue infiltrated with Xhg or water (Fig. 4). Some bacterial phytopathogens that infect the apoplast have evolved toxins that open stomata, increasing access to the leaf interior (reviewed in (20, 39)). Published works describe a model where Pst uses stomatal apertures as an entryway; using T3SS effectors to open stomata for bacterial entry and close the door behind them to create a water-soaked environment and reduce evaporation (19, 21, 22). Although *Xanthomonas* spp. and Pst both cause water soaking in the apoplast, we hypothesized that differences in stomata manipulation by each phytobacterial pathogen could impact bacterial entry to the apoplast and explain the differential outcomes of *S. enterica* persistence on infected leaves. We hypothesized that the closed stomata in Pst-infected tissue may prevent *S. enterica* entry when it arrives at 48 HPI, thus delaying *S. enterica* success in this environment. However, unlike published works in *A. thaliana* (21, 22), our data demonstrate that from 24 HPI to 72 HPI, Pst-infected tomato leaves maintain open stomata (Fig. 4). We predict that differences between our results here and published results (21, 22) are due to multiple differences in the investigated plant – microbe interactions. We anticipate that the primary driver of these disparities come from differences in the Pst strains used in the respective experiments. Pst DC3000 is pathogenic towards some tomato cultivars and *A. thaliana* while the Pst NY15125 strain used here is nonpathogenic towards *A. thaliana* (50). There are also known differences between these two bacterial strains in both T3SS effector profiles as well as virulence towards tomatoes, depending on the tomato host (30, 50). As T3SS effectors are vital for manipulating stomatal immunity, it is not surprising that profile differences could result in differential effects on stomata.

While leaves infected with either phytobacterial pathogens have open stomata when *S. enterica* arrives at 48 HPI, pathogen identity impacts stomatal response to *S. enterica* flagellin. Pst-infected leaves maintain open stomata in the presence of flagellin while Xhg-infected leaves initially close stomata upon flagellin induction and then reopen them within four hours (Fig. 4B). This short period of stomatal closure does not seem to impact *S. enterica* entry into the apoplast, as the droplet containing *S. enterica* is absorbed into leaf tissue in the same timeframe and *S. enterica* thrives in this environment (Fig. 1). While these results point to one difference between the two phytopathogens in terms of plant response to environmental cues, *S. enterica* has access to open stomata in both Xhg- and Pst-infected leaves. Thus, differential regulation of stomata does not explain why Pst fails to support *S. enterica* colonization as quickly as Xhg.

### Dispersal Within Host Tissue

For more than a century, plant pathologists have used disease severity indices to rate gross aspects of infection that are generally based on the percent area with visible symptoms (for review (51)). Yet, this disease quantification is a crude characterization of changes to the infected host and reveals nothing of the infected population’s current activities and dispersal since the bacterial pathogen is rarely isolated from diseased tissue. To quantify changes to the host following infection, [Leaf Lesion Detector](https://leaflesiondetector.streamlit.app/) (LLD), a new software application was created to quickly discriminate leaf tissue from background, identify color variation of leaves, measure the area of the leaf and area of discoloration within the leaf border, and calculate the discolored area relative to a fixed object (i.e. a U.S. dime). This application is publicly available and provides a new tool for researchers examining plant disease. Here, LLD was used to quantify disease progression in Xhg- and Pst-infected leaves (Fig. 2) and demonstrated that both phytopathogens produced quantitatively and qualitatively similar lesions, in abundance and size, over the course of infection. While no differences were noted that would explain the differential success of *S. enterica* in these infected plants, heterogenous patches of water soaking developed in both Xhg- and Pst-infiltrated leaves at 1 DPI (Fig. 3A). In contrast, infiltration immediately produces a homogenously water-congested area that quickly dissipates within several hours. The resulting heterogeneity of apoplast water soaking just one day later suggests that the phytopathogens may form microcolonies or that infection of host cells is not in synchronicity before water-soaking floods the entire infected area by 2 DPI (Fig. 3A). The formation of distinct subpopulations occurs in *P. syringae* pv. *phaseolicola* in the apoplast of bean leaves where bacteria are found clustered in microcolonies after infiltration (52). Multispecies interactions within these subpopulations even support the success and dispersal of non-pathogenic strains (52). Thus, although symptomology of Xhg- and Pst-infected leaves appears macroscopically similar, the infected and colonized apoplast reflects a more complex environment that requires further exploration to identify phytopathogen-specific mechanisms of niche establishment and host manipulation.

In terms of space, the carrying capacity of bacteria that colonize leaf surfaces have been well documented, at least for the phytopathogen Pst and the non-pathogenic *Pantoea agglomerans* (36, 53–59). On the leaf surface, bacterial aggregates are more likely to survive than individual cells, especially in unfavorable conditions on arrival sites (12). Despite this wealth of information, the carrying capacity of the apoplast remains poorly understood. We predict that the carrying capacity of the apoplast is dynamic and changes as the state of the host fundamentally transforms following infection. We and others have measured the size of the phytobacterial population within the apoplast by infiltration and subsequent enumeration at later times (60) (Fig. 3). Our results indicate that tomato leaves have similar carrying capacities for Pst and Xhg. Both phytopathogens reach plateaus at approximately the same level and time point in infection (Fig. 3). However, Xhg-infection enhances persistence of immigrating bacteria, such as *S. enterica,* while Pst-infection does not (Figs. 1 and 6). While these results indicate that space restriction doesn’t appear to be a primary mechanism for impacting immigrant success, it suggests that infection or resident status with the different genera results in unique niches within the same host.

Water congestion may also impact bacterial migration and dispersal within host tissues. Bacteria utilize swimming and other forms of motility to reach preferential niches, responding to environmental cues to reach essential nutrients (3, 15, 16, 61, 62). Here, we show that both Xhg and Pst remain susceptible to UV (Fig. 6), indicating that the phytopathogens spend more time on the leaf surface than *S. enterica,* even once water soaking has begun. Unlike *S. enterica,* Xhg and Pst may be transitioning in and out of the apoplast via stomata or through damaged tissue once lesions have developed. Bacterial phytopathogens use motility as a mode of dispersal within and between plants (12, 15, 36, 63). Thus, the observed UV susceptibility could reflect increased attempts of the phytopathogens to migrate to distal sites within or on the infected host. In contrast, our data suggest that, once within the apoplast, few *S. enterica* cells, if any, migrate back to the leaf surface. We also found that Pst infection appears to inhibit migration to the adjacent site for all arriving bacteria (*S. enterica,* Xhg, or Pst; Figs. 1 and 6). Previous results have suggested that Pst infection transforms the apoplast, either building physical barriers through biofilm production or manipulating the host response to prevent migration once inside the leaf tissue. For example, Pst infection in *A. thaliana* leads to restricted vascular flow to the infection site, which could also limit bacterial migration (64). Similarly, the ROS response to phytobacterial infection may link bacterial cells to cell wall components within the apoplast (65), inhibiting bacterial movement. A phytopathogen-specific mechanism during infection would explain why movement of immigrating bacteria is restricted in Pst-infected tissue while Xhg- infected tissue allows bacteria to move more freely.

### Type VI Secretion and Bacterial Competition

In searching for additional differences between Xhg and Pst that could explain the different impacts to *S. enterica* colonization, we focused on known virulence factors in the two phytopathogens. The genome sequences for both bacteria indicate that, while Pst contains factors for production of two Type VI secretion systems (T6SS), Xhg does not have the genes needed for this process. The T6SS is used by bacteria to deliver growth inhibitory toxins in a cell-cell contact dependent manner to competitor bacteria (for review (66)). Neither Pst T6SS is required for virulence in plants but both play a role in intermicrobial competition (67). Thus, we hypothesized that Pst uses its T6SSs to outcompete neighboring bacteria during colonization of the apoplast. If one or both Pst T6SSs inhibit or kill *S. enterica,* then the lack of these systems in Xhg could explain the success of *S. enterica* in Xhg-infected plants versus Pst-infected plants. Using an *in vitro* ‘killing assay’ for T6SS activity, we showed that *in planta* grown Pst inhibits growth of multiple bacteria, including *S. enterica,* Xhg, and even *in vitro* grown Pst (Fig. 5). These assay conditions mimic the arrival of naïve bacteria to an established Pst infection in leaves; bacterial strains are grown under the same conditions and mixed at the same ratios (1:1000; target strain to killer strain). Immigrating bacteria are susceptible to the T6SS toxin because they either don’t have the immunity protein to protect themselves (*S. enterica* and Xhg) or, because the previous growth conditions doesn’t trigger expression of the required immunity protein (Pst). In either case, the immigrating bacteria fall prey to the effects of the Pst T6SS toxin. As all immigrating bacteria, regardless of whether the immunity protein is present, eventually recover from the initial inhibition by Pst on leaves (Figs. 1 and 6), we suggest that Pst T6SS production is dynamic and only present during early time points in infection. Further experiments to directly address the role of the Pst T6SSs in *S. enterica* persistence will be needed to confirm the importance of one or both systems in this interaction.

To summarize, the leaf surface and apoplast present a complex and dynamic environment for bacterial communities, influenced by both host and bacterial factors. The fate of immigrant bacteria is shaped by their arrival conditions and the presence of established populations, which can create distinct niches and competition for resources. Nutrient availability, water congestion, water soaking, and stomatal responses play crucial roles in bacterial colonization and persistence. Finally, the presence of interbacterial competition factors, such as the Pst T6SS, further impacts colonization and survival. Further research is needed to dissect the molecular mechanisms underlying these processes and develop targeted strategies for disease control. A more comprehensive understanding of the bacterial infection and colonization mechanisms in plant tissues could ultimately inform crop protection strategies, enhance agricultural sustainability, and improve food safety.

## Materials and Methods

### Bacterial strains, media, and culture conditions

Strains used in this study are shown in Table 1. Kanamycin resistant Xhg was created by transforming Xhg 444 wildtype strain with pKTKan (68). Bacterial cultures were grown in lysogeny broth (LB) for *S. enterica*, nutrient broth (NB) for Xhg, and nutrient yeast extract dextrose broth (NYD) for Pst. All bacterial strains were incubated at 28°C with shaking at 200 rpm. The antibiotics nalidixic acid (Nal), kanamycin (Kan), and gentamicin (Gent) were used at concentrations of 20, 50, and 10 µg ml^−1^, respectively.

**Table 1.** List of Strains.

| Strain Designation | Genotype | Reference or source |
| --- | --- | --- |
| JDB1022 | *S. enterica* serovar Typhimurium 14028s; Kan<sup>R</sup> at the *attTn7* site | (2) |
| JDB1470 | *X. hortorum* pv. *gardneri* 444; Nal<sup>R</sup> | (2) |
| JDB1504 | *X. hortorum* pv. *gardneri* 444 + pKTKan; Kan<sup>R</sup> | This study |
| JDB1514 | *P. syringae* pv. *tomato* NY25; Kan<sup>R</sup> | Martin lab |
| JDB1515 | *P. syringae* pv. *tomato* NY25; Gent<sup>R</sup> | Martin lab |

### Plant inoculation

*Solanum lycopersicum* (tomato cultivar MoneyMaker) seeds were purchased commercially (Eden Brothers). This study complies with the relevant institutional, national, and international guidelines and legislation for experimental research on plants. Seedlings were cultivated in Professional Growing Mix (Sunshine Redi-earth) with a 16 h photoperiod at 24°C for five weeks. For colonization assays, Xhg and Pst bacterial cultures were grown for two days in NB or NYD, respectively, at 28°C, and *S. enterica* cultures were grown overnight in LB at 28°C. Bacterial strains were normalized to an optical density at 600 nm (OD_600_) of 0.2 (for *S. enterica* and Pst strains) and 0.3 (for Xhg) in sterile water. These OD_600_ values correspond to a bacterial population level of ∼10^8^ CFU/ml for the respective strains. <u>For dip inoculation</u>, normalized Xhg and Pst cultures were diluted 1:200 in sterile water for an inoculum level of ∼5x10^5^ CFU/ml. Prior to dip inoculation, 0.025% Sil-Wett was added to the bacterial inoculum. Tomato plants were dip-inoculated by inverting plants in the bacterial inoculum for 30 seconds with gentle agitation to prevent bacterial cell settlement. Plants were incubated at high humidity for 48 h in lidded, plastic bins under grow lights with a 12 h photoperiod at room temperature (∼26°C). After 48 h, plants were exposed to low humidity conditions (bin lids were removed) during the day and high humidity conditions (bin lids were replaced) during the night. <u>For infiltration experiments</u>, two leaflets on the third true leaf of MoneyMaker tomato plants were infiltrated with Xhg or Pst at ∼1x10^7^ CFU/ml, prepared as above for dip inoculation experiments, following published protocols (2, 69). Bacterial solutions were infiltrated into the abaxial leaf surface using a needleless syringe, and, for some experiments, infiltrated zones were delineated with permanent marker. Infiltrated plants were incubated in lidded, plastic bins as described above. For disease comparison of infiltrated plants, non-destructive images of leaflets were taken at multiple days post-infiltration using a Canon PowerShot ELPH 1901S camera.

### Image collection and processing

At multiple days post-dip inoculation, individual leaflets were removed, submerged in water for 10 seconds (to increase lesion visibility), gentle patted dry with kimwipes, and imaged for lesion quantification on a black velvet background using a Canon PowerShot ELPH 1901S camera. Images were cropped to center the leaflet and remove background. Leaf area and lesion analysis was performed with the custom image processing software: Leaf Lesion Detector v2023.2.0 (LLD), developed for this project. LLD automates manual image segmentation procedures (70). Results from LLD directly visually correspond to leaf lesions and also correlate with manually produced results (R^2^ = 0.95). The application is publically available at this url: https://leaf-lesion-detector.streamlit.app/. The code and the configuration settings used in this is work are also publically available on GitHub (pending publication): https://github.com/UW-Madison-DSI/plant-pathology-image-processor.

### Immigrant Arrival

*S. enterica*, Xhg, and Pst cultures were grown as described above, normalized to OD_600_ = 0.1, and diluted in sterile water for a final concentration of ∼10^6^ CFU/ml. Normalized and diluted immigrant cultures were applied as 15 μl droplets to the adaxial surface of infiltrated leaf tissue, which included leaflets infected with Xhg or Pst for 2 DPI, as well as leaflets freshly infiltrated with sterile water, as done previously (3). The droplet “arrival site” was denoted with permanent marker and is depicted in Figure 1 for clarity. Inoculated plants were incubated at room temperature for 3 h to allow for droplet absorption and then moved under grow lights until sampling. Bacterial inoculum were diluted and plated for population counts.

### Bacterial population sampling

Bacterial populations on leaves were determined as described (2, 3, 69). Briefly, at indicated times, two individual leaflets were removed from each plant, and the adaxial surface of one leaflet per plant was treated with 254 nm UV radiation (Stratalinker UV Crosslinker 1800) at 150,000 μJ/cm^2^ (for *S. enterica*) or 300,000 μJ/cm^2^ (for Xhg or Pst). Irradiated leaflets were chosen at random for each plant. Two 79 cm^2^ leaf discs were taken from each leaflet using a surface sterilized cork borer. One disc was removed from site of droplet application and designated as the arrival site (Fig. 1). The other disc was sampled at an adjacent, nonoverlapping region within the infiltration zone and termed the adjacent site (Fig. 1). Samples from four plants per treatment per time point were individually homogenized in 500 μl of sterile water in microfuge tubes using a 4.8 V rotary tool (Dremel, Mt. Prospect, IL) with microcentrifuge tube sample pestle attachment (Fisher Scientific). Homogenates were diluted and plated to enumerate bacteria on LB Kan (for *S. enterica*), NB Nal or NB Kan (for Xhg), or NYD Kan or Gent (for Pst) agar plates. Remaining leaf homogenate was incubated with media amended with kanamycin to enrich for immigrating bacteria below the limit of detection in the above plating method (3). Enrichments were plated on media with kanamycin after 1 day (*S. enterica*) or 2-3 day (Xhg and Pst) incubation at 28°C. Resulting colonies were counted after overnight incubation at 37°C (for *S. enterica*) or after incubation for 2-3 days at 28°C (for Xhg and Pst) to determine bacterial populations. Experiments were performed with three biological replicates.

### Electrolyte leakage

Plants were infiltrated with Xhg, Pst, or water alone, as described above. Leaf tissue samples were collected at 0, 1, 2, 3, and 4 DPI for bacterial populations and electrolyte leakage measurements. Four leaf discs were taken from the infiltrated zones of leaflets from the third true leaf of each plant. One disc was used to enumerate bacterial populations (as described above), and three discs were pooled and quantified for changes in electrolyte leakage (conductance) as previously described (71, 72). Briefly, leaf discs were placed in a 12-well plate containing 4 ml sterile water per well, three leaf discs per well. Leaf discs were washed for 30 minutes with gentle agitation, and the water was removed and replaced with 4 ml fresh, sterile water. Baseline conductance values from each well were immediately measured using an ECTestr11 + MultiRange electrical conductance probe. Plates were incubated under grow lights for 6 h, and final conductance values were collected from each well. Data are expressed as differences between baseline and final conductance readings as a proxy for cellular damage and electrolyte leakage. Experiments were performed with three biological replicates.

### Stomatal aperture imaging and analysis

Plants were infiltrated with Xhg, Pst, or water alone, and stomatal apertures were measured using a dental resin technique (73, 74). Bacterial suspensions were prepared as described above and infiltrated as ∼1 cm^2^ areas on leaflets from the third true leaf. To capture stomatal apertures at 1, 4, 24, 48, and 72 HPI, impressions of the leaf surface were taken using light-body vinylpolysiloxane (VPS) dental resin. Resin was applied in a thin layer to the adaxial surface of leaves over the infiltrated area, dried for ∼5 min, and then removed. Resin impressions were coated with clear nail polish. Once dried and removed, the nail polish mirrors the leaf surface impression. Nail polish imprints were placed on cover slips, and stomata were imaged with a Cytation 7 Cell Imaging Multi-Mode Reader (Biotek). Infiltrated areas on leaflets were outlined with permanent marker, which transfers to both the resin and nail polish and allows for identification of infiltrated tissue under the microscope. Stomatal apertures were measured within infiltrated areas using ImageJ (75) (n > 100 stomata per treatment), and resulting length to width ratios are presented in Fig. 4. To examine stomatal responses to flagellin, leaflets were infiltrated with Xhg, Pst, or water, as described above. At 48 HPI, purified flagellin (50ng/ml; Fisher Scientific) was applied as a droplet to the adaxial surface of the infiltrated areas. Resin impressions were taken at 1 or 4 HPA, and stomatal apertures were measured as described above. Experiments were performed with three biological replicates.

### Bacterial growth inhibition assays (“killing” assays)

Plants were infiltrated with Xhg, Pst, or water alone, as described above. At 48 HPI, four 79 cm^2^ leaf discs were taken from infiltrated areas using a surface sterilized cork borer, pooled, and homogenized in 500 μl of sterile water. This procedure was repeated for each treatment to collect concentrated homogenates to act as the “killer” suspensions. Target strains were prepared as described above for plant inoculations except that strains were normalized to OD_600_=0.5. Normalized target strains were mixed with concentrated killer strain homogenates at a ratio of 1:1000. Killer-target mixtures, as well as killer and target strains alone, were spotted to agar plates (LB for *S. enterica*, NB for Xhg, or NYD for Pst) and incubated at 28°C for 24 h. Agar cores were then excised from plates and vortexed thoroughly in 1 ml sterile water to collect bacterial cells. Suspensions were diluted and plated for bacterial enumeration. Resulting colonies were counted after overnight incubation at 37°C (for *S. enterica*) or after incubation for 2-3 days at 28°C (for Xhg and Pst) to determine bacterial populations. Experiments were performed with three biological replicates.

### Statistical Analysis

All statistical analyses were performed using R software (version 2.14.1; R Development Core Team, R Foundation for Statistical Computing, Vienna, Austria [http://www.R-project.org]) as described (76). Three biological replicates were performed for each experiment. The ANOVA and Tukey’s HSD test were used to compare treatment at each time point. The one sample t-test was used to compare bacterial populations to the initial arriving populations on tomato leaves. Results were considered statistically significant at *P* < 0.05.

## ACKNOWLEDGEMENTS

Funding was provided by the Food Research Institute, University of Wisconsin-Madison. The funders had no role in study design, data collection and interpretation, or the decision to submit the work for publication. We would also like to thank G. Martin (Cornell Univ.) for marked Pst NY25 strains and acknowledge JDB’s lab members for helpful discussions.

KNC and JDB designed the experiments for this work, KNC, EGG, DN and SCZ performed the experiments, KNC and JDB analyzed and interpreted the data, and ASI and IM designed the LLD app. KNC and JDB wrote the main manuscript text, and all authors reviewed the manuscript.

